# Human *SYNGAP1* Regulates the Development of Neuronal Activity by Controlling Dendritic and Synaptic Maturation

**DOI:** 10.1101/2020.06.01.127613

**Authors:** Nerea Llamosas, Vineet Arora, Ridhima Vij, Murat Kilinc, Lukasz Bijoch, Camilo Rojas, Adrian Reich, BanuPriya Sridharan, Erik Willems, David R. Piper, Louis Scampavia, Timothy P. Spicer, Courtney A. Miller, J. Lloyd Holder, Gavin Rumbaugh

**Author notes:** GSK 1250 S Collegeville Rd., Collegeville, PA 19426. **Corresponding Author:** Gavin Rumbaugh, Department of Neuroscience, 130 Scripps Way #3B3, Jupiter, FL 33458.

## Abstract

*SYNGAP1* is a major genetic risk factor for global developmental delay, autism spectrum disorder, and epileptic encephalopathy. *De novo* loss-of-function variants in this gene cause a neurodevelopmental disorder defined by cognitive impairment, social-communication disorder, and early-onset seizures. Cell biological studies in mouse and rat neurons have shown that *Syngap1* regulates developing excitatory synapse structure and function, with loss-of-function variants driving formation of larger dendritic spines and stronger glutamatergic transmission. However, studies to date have been limited to mouse and rat neurons. Therefore, it remains unknown how *SYNGAP1* loss-of-function impacts the development and function of human neurons. To address this, we employed CRISPR/Cas9 technology to ablate *SYNGAP1* protein expression in neurons derived from a human induced pluripotent stem cell line (hiPSC). Reducing SynGAP protein expression in developing hiPSC-derived neurons enhanced dendritic morphogenesis, leading to larger neurons compared to those derived from isogenic controls. Consistent with larger dendritic fields, we also observed a greater number of morphologically defined excitatory synapses in cultures containing these neurons. Moreover, neurons with reduced SynGAP protein had stronger excitatory synapses and expressed synaptic activity earlier in development. Finally, distributed network spiking activity appeared earlier, was substantially elevated, and exhibited greater bursting behavior in *SYNGAP1* null neurons. We conclude that *SYNGAP1* regulates the postmitotic maturation of human neurons made from hiPSCs, which influences how activity develops within nascent neural networks. Alterations to this fundamental neurodevelopmental process may contribute to the etiology of *SYNGAP1*-related disorders.

## Introduction

Pathogenic loss-of-function variants in the *SYNGAP1* gene are causally-linked to a range of neuropsychiatric disorders, including global developmental delay (GDD)/intellectual disability (ID) (Hamdan et al., 2009; Rauch et al., 2012; Deciphering Developmental Disorders, 2015, 2017) and severe epilepsy (Carvill et al., 2013; von Stulpnagel et al., 2015; Vlaskamp et al., 2019). *SYNGAP1* is also strongly implicated in autism spectrum disorders (Rauch et al., 2012; O’Roak et al., 2014) and was recently identified as one of three genes that impart the highest risk for developing autistic features (Satterstrom et al., 2020). While pathogenic variants in *SYNAGP1* are overall rare, they are common relative to the pool of genes capable of causing sporadic neurodevelopmental disorders, explaining up to ~1% of GDD/ID cases (Berryer et al., 2013; Parker et al., 2015). While the exact incidence of *SYNGAP1* pathogenicity remains unknown, early estimates are 1/10,000 (Parker et al., 2015; Weldon et al., 2018), which is in the range of other monogenic disorders that are more extensively studied by the scientific community. Causality of *SYNGAP1* pathogenicity is now established because there are no known loss-of-function variants in >141,000 neurotypical individuals from the gnomAD database and all known patients with clearly pathogenic variants are diagnosed with a neurodevelopmental disorder (Mignot et al., 2016). Moreover, the pLI ratio for *SYNGAP1* is 1 (Lek et al., 2016; Jimenez-Gomez et al., 2019), demonstrating that the human gene is extremely intolerant of loss-of-function variants. Based on substantial clinical evidence, proper *SYNGAP1* expression is required for normal human brain development and function.

*Syngap1* gene function has been studied in rodent neurons (Kilinc et al., 2018; Gamache et al., 2020). *Syngap1* is a potent regulator of dynamic processes required for Hebbian plasticity at excitatory synapses. Heterozygous knockout mice exhibit deficits in hippocampal LTP evoked through a variety of synaptic stimulation protocols (Komiyama et al., 2002; Kim et al., 2003). This function of SynGAP protein is consistent with cognitive impairment commonly observed in *SYNGAP1* patients because Hebbian plasticity at excitatory synapses is thought to contribute importantly to learning. Genetic re-expression of *Syngap1* in adult mutant mice rescues hippocampal LTP and associated downstream signaling pathways (Ozkan et al., 2014). Thus, SynGAP regulation of synapse plasticity is a dynamic function of the protein that is retained throughout life. Hundreds of genes regulate synaptic plasticity as referenced by the Gene Ontology Browser (366 genes; http://www.informatics.jax.org/vocab/gene_ontology/GO:0048167). However, most of them do not cause disease when heterozygously expressed, as is the case for *SYNGAP1* (Carvill et al., 2013; Deciphering Developmental Disorders, 2015, 2017; Satterstrom et al., 2020). Therefore, *SYNGAP1* likely has additional functions beyond regulation of synapse plasticity that contribute to disease etiology. Indeed, there are additional reported functions of the *Syngap1* gene. SynGAP expression in developing mouse neurons acts to regulate the maturation rate of excitatory synapse strength and this function is independent from its role in plasticity. SynGAP protein expression rises quickly during postnatal development (Gou et al., 2020) and its expression during this period is critical for shaping the strength of nascent excitatory synapses (Clement et al., 2012; Clement et al., 2013). *Syngap1* heterozygous mice have enhanced excitatory synapse function in the developing cortex and hippocampus, which is thought to contribute to early onset of behavioral deficits and seizures observed in these animals. In contrast to Hebbian processes, this function of rodent *Syngap1* is linked to biological process unique to developing neurons. Enhanced baseline excitatory synaptic strength in hippocampal neurons is transiently observed during the first three postnatal weeks of brain development and inducing heterozygosity of *Syngap1* beyond this period has minimal effect on resting synaptic function in these neurons (Clement et al., 2012).

The understanding of how this gene contributes to disease-relevant biology is limited because information on its function in human neurons is lacking. This is limiting because there are fundamental differences in how human and rodent brains develop. For example, humans express neoteny, or slowing of development, which is thought to promote an extended period of neural network refinement that promotes higher cognitive functions. An example of neoteny at the neurobiological level is the relative pace of human neuron development compared to rodents (Petanjek et al., 2011; Charrier et al., 2012), with human neurons exhibiting a much slower pace of postmitotic differentiation. Given that *Syngap1* alters measures of neuronal maturation in rodents (Clement et al., 2012; Clement et al., 2013; Aceti et al., 2015), this function of the gene may be amplified in slower developing human neurons. To test this idea, we created *SYNGAP1* knockout human induced pluripotent stem cell (hiPSC) lines using CRISPR/Cas9 technology. These iPSCs were then differentiated into neurons (iNeurons) and cultures were assessed for various parameters of neuronal maturation. We found that human iNeurons lacking SynGAP expression exhibited accelerated dendritic morphogenesis, increased accumulation of postsynaptic markers, early expression of synapse activity, enhanced excitatory synaptic strength, and early onset of neural network activity. We conclude that *SYNGAP1* regulates the postmitotic differentiation rate of developing human neurons and disrupting this process impacts the function of nascent neural networks. These observations in human neurons are consistent with findings from rodent studies, indicating that control of neuronal maturation is a species-conserved function of the gene. Therefore, disruptions to this fundamental neurodevelopmental process may contribute to the etiology of *SYNGAP1*-related brain disorders.

## Material and Methods

### Maintenance of hiPSC cultures

All hiPSC work was performed in accordance with approved protocols from appropriate Institutional Review Boards. All products were purchased from Thermo Fisher Scientific unless otherwise noted. The stable human episomal Cas9 hiPSC cell line was obtained from Thermo Fisher Scientific (A33124) and was expanded according to the manufacturer’s suggested protocol. This line was previously used for generating neurons (Sridharan et al., 2019). Briefly, culture plates were coated with Vitronectin-N (A14700), diluted 1:100 in DPBS (14190094), and incubated at 37 °C for at least 1 h prior to iPSC plating. Cryopreserved iPSC cells were gently thawed in a 37 °C water bath and transferred to a 15 mL conical tube with Complete iPSC Medium + 1% RevitaCell supplement (A2644501). Cells were then centrifuged at 200 x g for 4 min and the iPSC pellet was re-suspended in fresh medium and plated on vitronectin coated flasks. Twenty-four hours later, cells were switched and maintained in Complete Stemflex Medium (w/o RevitaCell) with daily medium changes until 70% confluent. Cells were then harvested with TrypLE Select (12563011) and further maintained or plated for experimental purposes. For limiting dilution cloning, iPSCs were plated in 96-well plates coated with 2.5μm/ml rhLaminin-521 (A29248).

### Generation of *SYNGAP1* KO hiPSC lines

Guide RNA (*g*RNA) sequences targeting exon 7 of *SYNGAP1* were selected using the Zhang lab CRISPR design tool (http://zlab.bio/guidedesign-resources) and acquired from IDT in single guide RNAs (*sg*RNA) format. Cas9-iPSCs were transfected with *sg*RNAs by using Lipofectamine CRISPRMAX (Thermo Scientific,CMAX00001) according to manufacturer’s instructions. Editing efficiency of individual sgRNAs was determined using GeneArt Genomic Cleavage Detection Kit (Thermo Scientific, A24372). *sg*RNA-5 (target sequence 5’-TCTTTCGGCCGCAGACCGAC-3’) demonstrated the highest efficiency and was selected for downstream applications. To generate the *SYNGAP1* KO iPSC lines, cells were transfected with *sg*RNA-5. Twenty-four hours after the transfection, cells were plated in rhLaminin-521 coated 96-well plates with an average density of 0.5 cells/well. Colonies derived from a single cell were expanded and cryopreserved with *Recovery* cell culture freezing medium (Thermo Scientific, 12648010). Approximately 70 colonies originating from a single cell were analyzed for Indels around the *sg*RNA targeting site. Multiple clones with either unedited (WT) or edited (potential KO) sequences were isolated and expanded. Potential KO clones with “clean” Sanger sequence traces were prioritized. Pluripotency of individual clones were confirmed via TaqMan Array Human Stem Cell Pluripotency Panel (4385344) according to manufacturers’ instructions. Each expanded clone was tested, and confirmed negative, for mycoplasma contamination using Universal Mycoplasma Detection Kit (ATCC, 30-1012K).

### Whole exome sequencing (WES)

Genomic DNA from the four experimental clones and a sample of the original Cas9 iPSC line (before CRISPR transfection) were extracted using PureLink Genomic DNA mini Kit (Invitrogen #k1820-02) using included instructions. Genomic DNA from each of the five samples was shipped to HudsonAlpha Institute for Biotechnology, Genome Sequencing Center (Huntsvile, AL) for WES.

#### Library Preparation and Quality Control

DNA samples were normalized to 1,000ng of DNA in 50μl of water. Following normalization, samples were acoustically sheared via Covaris LE-220 instrument to a final fragment size of ~350-400bp. The sheared DNA was then transformed into a standard Illumina paired-end sequencing library via standard methods. The sheared DNA was end-repaired and A-tailed using Roche-Kapa End-Repair and A-Tailing kits under the manufacturer’s recommended conditions. Standard Illumina paired-end adaptors were ligated to the A-tailed DNA. Following ligation, the reactions were purified using AMPure XP beads. The purified ligated DNA was amplified via PCR using Roche KAPA HIFI PCR reagents using 4 cycles of PCR. The primers used in the PCR step introduced 8-base, unique, dual indexes in the i5 and i7 positions to allow sample identification/demultiplexing following sequencing. The final library was quality controlled using size verification via PerkinElmer LabChip GX and real-time PCR using the Roche KAPA SYBR FAST qPCR Master Mix, primers and standards according to the manufacturer’s directions. Libraries were normalized to 1.4 nM stocks for use in clustering and sequencing.

#### IDT Exome Capture and Quality Control

Post-library construction, samples were multiplexed for capture at 5 samples per pool with each sample contributing a maximum of 300ng or a minimum of 200ng to each pool. Pooled samples were purified with beads and eluted in a volume of 30μl. Pooled samples were hybridized with the NimbleGen SeqCap EZ Exome v3 probes with minor modifications for automation. Briefly, multiplexed samples were dried down in the presence of COT-1 and a blocker mix for 1.5 hrs. Libraries were then resuspended in a mix of hybridization buffer and baits. Libraries were hybridized overnight at 65°C (72 hrs). Post-hybridization takedown occurred 72 hours later. Briefly, captured libraries were bound to streptavidin beads. Once bound, washing occurred per manufacturer’s recommendations. Final elution of captured libraries was in 20μl of nuclease-free water. Libraries were amplified with six cycles of PCR and a final bead purification. Post-hybridization exome concentrations were measured via Picogreen and library sizes were determined via the LabChip GX Touch HT (PerkinElmer). Additionally, libraries were quantitated with real-time PCR using the KAPA Library Quantification Kit (Roche) per manufacturer’s instructions to determine final library nanomolarity. Final exome libraries were pooled at a concentration of 1.8nM. The pooled exome libraries were distributed across four lanes on an S4 flow cell and sequenced using 150 base pair paired-end approach on a NovaSeq 6000 instrument (Illumina). All sequencing was performed on the Illumina NovaSeq 6000 platform by loading a pool samples to the equivalent loading of 24 samples per flowcell. Following sequencing all basecalling was performed using standard Illumina software to generate the final FASTQ files for each sample. Alignment and variant calling was performed via the Edico/Illumina DRAGEN pipeline to verify coverage and performance. Samples yielded a minimum of 440M paired reads at 150nt read length with a mean coverage of greater than 30X.

### Karyotyping

Karyotyping was performed as previously described (Sridharan et al., 2019). Briefly, differentiated iNeurons were assessed for any chromosomal aberrations using the Karyostat™ assay (Thermo Fisher).

### Generation of induced Neurons (iNeurons) from Cas9-iPSC single cell clones

Ngn2 transcription factor induced iNeurons were generated as previously described (Sridharan et al., 2019) with minor modifications. Briefly, Cas9-hiPSCs were harvested using TrypLE Select and 2 million cells were plated on vitronectin coated T75 flask on day 1. On day 2, medium was removed, and an appropriate amount of lentivirus expressing Ngn2 (Addgene, 52047) and rtTA (Addgene, 20342) were administered in Complete Stemflex Medium including 1% RevitaCell (MOI 2 for both lentivirus). After 24 h, the medium was aspirated and replaced with Induction Media induce TetO gene expression. The next day, medium was refreshed with Induction Media + 2 μg/mL Puromycin (A1113803) which was included for selection of iNs. Twenty-four hours later, iNeurons were harvested using Accutase (A1110501) and plated on PDL coated plates in iNeuron Maintenance Media (Neurobasal (211103049) + 1% GlutaMax (35050061) + 2% B27 (17504044) + 10ng/ml BDNF (PHC7074) + 10ng/ml GDNF (PHC7036) + 10ng/ml NT3 (PHC7045) + 2.5% FBS (10082139), all from Thermo) + 10 μg/ml FuDR (Sigma, F0503) along with primary rat glia (neuron/glia ratio 2.5/1). Half of the medium was changed with fresh iNeuron Maintenance Media every 4-5 days.

### Dendritic Tracing

Each well of a 96-well imaging plate contained ~50,000 cells per well, consisting of ~32,000 human induced Neurons (iNs) + ~18,000 Primary Rat astrocytes along with 0.1% (~ 50 per well) of EGFP positive human induced Neurons derived from the same clone. eGFP-positive iNeurons were created through a separate induction as stated above, except that an additional lentivirus expressing eGFP under the control of a TET-responsive promoter was included (Addgene Cat # 30130). eGFP-positive neurons were mixed with eGFP-negative neurons in the 96-well plates. iNeurons derived from either of the WT or KO clones were compared by tracing primary (originating from the soma), secondary, and tertiary dendrites, as well as total dendrite length. Tracing data was obtained by imaging live iNeurons at DIV45 with an InCell Analyzer 6000 automated confocal microscope (20X magnification). A sample of 30 randomly selected neurons per genotype (n= 3 per well x 10 wells in a 96-well plate) was selected and then dendrites were traced with the Simple Neurite Tracer (SNT) software plugin distributed by Fiji-ImageJ. Data represents the average lengths in microns for all subtypes of dendrites.

### Immunocytochemistry

iNeurons were re-plated, along with primary rat astrocytes, at a density of iNeurons 200,000/120,000 astrocytes per well, on 15mm cover glass coated with PDL/Fibronectin (Neuvitro, GG-15-Fibronectin), in 24-well plates. At DIV30-45, cells were fixed and labelled with primary antibodies: anti-PSD95 (mouse-raised; Abcam Cat# ab2723-100ug), anti-GluA1 (Rabbit-raised; Cell Signaling Technology Cat# 13185S) and anti-MAP2 (Guinea pig-raised; Synaptic Systems Cat # 188004). Then, secondary antibodies were applied (Goat anti-mouse Alexa 488, Abcam Cat# ab150113-500ug, Goat anti-rabbit Alexa 568, ThermoFisher Cat# A11036, Goat anti-guinea pig Alexa 647 ThermoFisher Cat# A21450). Images of neurons from multiple coverslips per culture were taken under UPlanSApo 100× 1.4 NA oil-immersion objective mounted on Olympus FV1000 laser-scanning confocal microscope (1 image = 1 field-of-view). Neuronal somas from individual fields-of-view were manually calculated based on raw MAP2-signals. Total area of MAP2/field-of-view was determined on the area of mask of MAP2 signal. Number of detected particles of GLUA1 and PSD95 per field-of-view was determined based on threshold-based signal masks. Thresholds were kept constant across all images.

### Immunoblotting

iNeurons were co-cultured with rat glia (500,000 induced neurons, 100,000 glia) seeded on 12-well plates. After 30-60 days in culture, media was removed and the wells were washed with PBS, after which the PBS was replaced with 200ul of RIPA buffer (Cell Signaling Technology, Danvers, MA) containing Phosphatase Inhibitor Cocktails 2 and 3 (Sigma-Aldrich, St. Louis, MO) and MiniComplete Protease Inhibitor Cocktail (Roche Diagnostics), the wells were scraped using a sterile cell scraper on each well and transferred to tubes in dry ice, and stored at −80°C. After thawing on ice, samples were sonicated using a probe sonicator 5 times with 2 sec pulses. Sample protein levels were measured (Pierce BCA Protein Assay Kit, Thermo Scientific, Rockford, IL), and volumes were adjusted to normalize microgram per microliter protein content. 10 μg of protein per sample were loaded and separated using SDS-PAGE on 4-15% gradient stain-free tris-glycine gels (Mini Protean TGX, BioRad, Hercules, CA), transferred to low fluorescence PVDF membranes (45 μm) with the Power Blotter-Semi-dry Transfer System (ThermoFisher Scientific). Membranes were blocked with 5% powdered milk in buffer and probed with pan-SynGAP (1:1,000, #5539, Cell Signaling) or SynGAP-α2 (abcam, ab77235), overnight at 4°C and HRP-conjugated anti-rabbit antibody (1:2,000, W4011, Promega) for 1 hr at room temperature followed by ECL signal amplification and chemiluminescence detection (SuperSignal West Pico Chemiluminescent Substrate; Thermo Scientific, Rockford, IL). Blot band densities were obtained using the ChemiDoc imaging system (BioRad). SynGAP levels of immunoreactivity were assessed by densitometric analysis of generated images with ImageJ. Values were normalized to total protein levels obtained from blots prior to antibody incubations.

### Whole Cell Electrophysiology

iNeuron measurements were performed up to DIV50 as described with minor modifications (Sridharan et al., 2019). For current studies, iNeurons were co-cultured with cryo-recovered primary rat astrocytes (seeded at 20,000 iNs + 10,000 astrocytes per well) in 24-well plate on 15 mm coverslips. The co-cultures were maintained in plating medium and additionally supplemented with 10 μg/mL FUDR (cat. no. F0503, Sigma). For whole cell recordings, intrinsic electrical properties were inspected immediately after gaining access to the cell and miniature excitatory synaptic currents were recorded in the presence of TTX (0.5 μm) at room temperature in voltage-clamp configuration (cells were held at −60 mV with a Multiclamp 700B amplifier, Molecular Devices). The bath solution contained (in mM): 140 NaCl, 5 KCl, 2 CaCl_2_, 2 MgCl_2_, 10 HEPES-NaOH and 10 Glucose (pH to 7.4 adjusted with NaOH). Pipettes pulled from borosilicate glass capillary tubes (cat. no. G85150T-4, Warner Instruments) using a P-97 pipette puller (Sutter Instrument) were filled with the following intracellular solution (in mM): 123 K-gluconate,10 KCl, 1 MgCl_2_, 10 HEPES-KOH, 1 EGTA, 0.1 CaCl_2_, 1 K_2_-ATP, 0.2 Na_4_-GTP and 4 glucose (pH adjusted to 7.4 with KOH). Resistance of the pipettes filled with the intracellular solution was between 3-5 mΩ. Series resistance was monitored without compensation with 5 mV depolarizing steps (200 ms) induced every 60s to ensure sufficient and stable electrical access to the cell. Data were sampled at 10 kHz, post hoc filtered and analyzed offline using Clampfit (Molecular Devices). Single peak mEPSCs were detected using a semiautomated sliding template detection procedure in which the template was generated by averaging multiple spontaneous currents. Each detected event was visually inspected and discarded if the amplitude was <7 pA.

### Microelectrode Array (MEA) Analysis

#### Cell culture and NPC differentiation

Individual *SYNGAP1* WT and KO iPSC clones were maintained on Matrigel-coated plates in Stem Flex media (Fisher Scientific). Neural Progenitor Cells (NPCs) were differentiated from iPSCs using a dual SMAD inhibition protocol (Jiang et al., 2017). Briefly, stem cell lines were dissociated using Accutase and embryoid bodies were generated from the stem cell lines in the Aggrewells using Neural Proliferation medium (NPM) along with BMP and WNT inhibitors (Dorsomorphin: DM; 4mM and SB-431542:SB 10mM; Sigma Aldrich), administered on Day 2 of neural induction. At ~ Day 5, EBs were gently collected and plated on Matrigel coated plates for the formation of rosettes. To promote dorsalization, 10mM Cyclopamine (CCP; Stem Cell technologies) was added to the plates starting Day 6. Both inhibitors and CCP were added to the media until ~ Day 9. Rosettes were collected between Day 14 and 16 and plated on gelatin-coated plates so that the non-neural cells were preferentially removed from floating neural progenitors, which were then dissociated to form a monolayer culture of neural progenitor cells. NPCs were grown and expanded on Matrigel coated plates before the cells were plated directly on a MEA plate for neuronal differentiation.

#### MEA analysis and neuronal differentiation

We employed an MEA system (Axion Biosystems) to perform neurophysiological characterization of iNeurons. Neuronal differentiation of NPCs was performed directly on MEA plates. 1.6×10^4^ NPCs suspended in a 5μl droplet of NPM (neural precursor medium) were plated as on top of a 16-electrode array (area ~1mm^2^) inside a single well of 48-well MEA plate pre-treated with 0.1% PEI solution prepared in borate buffer (pH=8.4). Two days later, neuronal differentiation was initiated using in neuronal induction medium (NIM, prepared from equal volumes of DMEM/F12 and neurobasal medium without growth factors) prepared in-house. NIM was exchanged every other day for 7 days. Differentiation of the NPCs into forebrain cortical neurons was performed using previously established neuronal differentiation medium, NDM, which includes a cocktail of differentiation factors (BDNF, GDNF, NT-3, dibutyryl-cAMP, ascorbic acid) (Jiang et al, 2017 Nature Comm). Post-differentiation, NDM was replaced with BrainPhys for further maturation (Stem Cell technologies), and neurons were cultured for at least one week before neuronal activity was recorded. Neuronal activity was recorded continuously for 5 minutes from the multi-well MEA plate each week until 6 weeks of neuronal maturation, post-differentiation. Field potential changes were recorded and analyzed using Axis Navigator and Axis metric plotting software (Axion Biosystems). Temporal raster plots were generated using Neural Metric Tool software (Axion Biosystems). For data analysis, a burst was identified as a group of at least 5 spikes, separated by an inter-spike interval (ISI) of less than 100 milliseconds. Network bursts were defined as a minimum of 50 spikes with a maximum ISI of 100 ms covering at least 35% of electrodes in a well.

### Statistics

GraphPad Prism 8 software was used for all statistical analysis. All data were tested for normality. Accordingly, parametric or non-parametric tests were applied. For tracing data analyses, clonal comparisons were performed using Kruskal-Wallis test followed by Dunn’s multiple comparison test. For genotype comparisons Mann-Whitney tests were applied. For immunostaining experiments, Mann-Whitney tests or unpaired two-tailed t-tests were used. For clonal comparisons of electrophysiological data Kruskal-Wallis followed by corrected Dunn’s multiple comparison tests or One-way ANOVA followed by Tukey tests were used. Statistical differences of percentage mEPSC-expressing neurons were determined by Fischer exact test pair-wise comparisons. For genotypic comparisons of whole cell electrophysiological data, Mann-Whitney U tests or Unpaired t-tests were performed. When comparing cumulative probability data between clones or genotypes the Kolmogorov-Smirnov test was used. For multielectrode array studies, statistical analyses among clones was performed using two-way RM ANOVA followed by Tukey’s multiple comparison test and for genotype comparisons two-way RM ANOVA followed by Bonferroni’s multiple comparison. Data throughout the text are presented either as box-and-whisker plots where the center, boxes and whiskers represent the median, interquartile range, and min to max, respectively, or as mean ± SEM. Differences were considered to be significant for p < 0.05. Exact p-values are reported when provided by the software.

## Results

To create *SYNGAP1* null hiPSCs, we performed CRISPR editing of a common exon within the human locus. Exon 7 was targeted **(Fig. 1A)** for non-homologous end joining (NHEJ) repair for the following reasons: 1) it is a common exon present in most, if not all, *SYNGAP1* transcripts (McMahon et al., 2012; Gou et al., 2020); 2) it is downstream of multiple stop-gain or small indel patient-specific variants (Jimenez-Gomez et al., 2019); 3) targeting it in other species results in ablation of SynGAP protein (Kim et al., 2003; Clement et al., 2012). Four single cell clones were identified and selected for downstream analysis. These clones contained either an edited (KO - Clone #4 and #38) or unedited (WT-Clone #6 and #30) Exon 7 **(Fig. 1A-B)**. Sanger sequencing of the putative KO clones contained “clean” sequence both up and downstream of the Cas9 cut site. Importantly, karyotyping analysis **(Fig. 1C)** revealed no large alterations to chromosomal structure in any of the four clones. Moreover, each of the clones passed self-renewal and pluripotency checks and tested negative for mycoplasma contamination. The type of variants present, combined with the Sanger sequence traces, suggested that both clones contained bi-allelic indel frameshift variants, which would be expected to cause nonsense mediated decay of *SYNGAP1* transcripts and disruption to SynGAP protein. To test this prediction, glutamatergic neurons were produced from each of the four clones using the Ngn2 induction method (Zhang et al., 2013). After ~30-60 days of neuronal development, samples were immunoblotted for SynGAP protein levels. As predicted, neurons derived from both “KO” clones had significantly lower levels of SynGAP protein than “WT” clones. Reduced SynGAP signal was observed with antibodies recognizing either a core region of the protein (Pan-SynGAP), or to the C-terminus of a specific splice variant (α2; **Fig. 1D-E**). Given that SynGAP signal is ~10% of control levels, the two KO clones appeared to produce iNeurons with nominal SynGAP protein expression.

**Figure 1.**
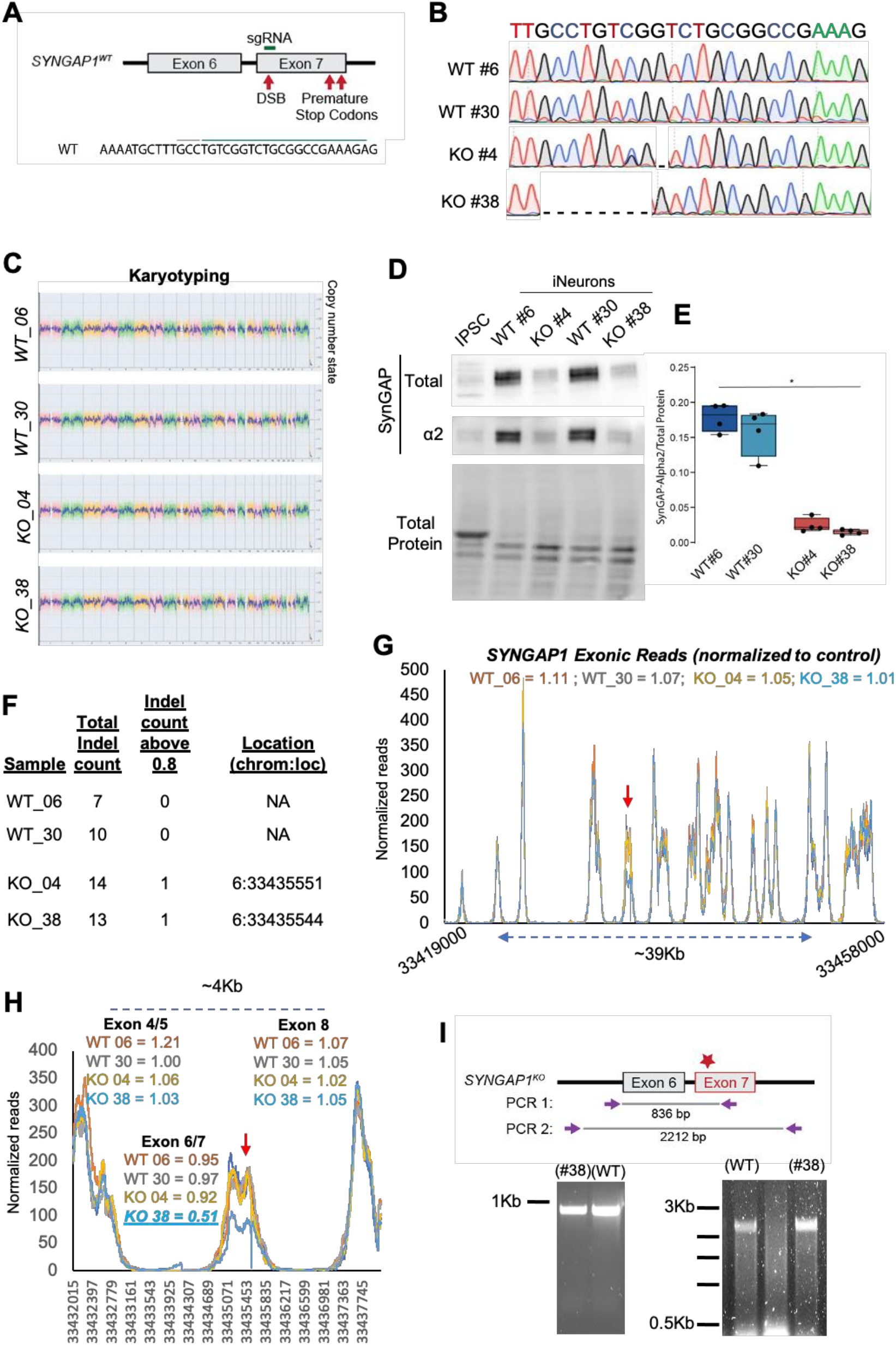
Development of isogenic *SYNGAP1* knockout hiPSCs. **(A**) Cartoon showing clone-specific mutations in the *SYNGAP1* gene. **(B)** Sanger sequencing for one WT clone and two *SYNGAP1* mutant clones derived from the CRISPR experiment. (**C**) Whole genome view of iNeurons from WT#6, WT#30, KO#4 and KO#38 clones depicting a copy number value of 2 cross all chromosomes (expect for the Y-chromosome which is not detected) revealing normal (female) karyotype with no chromosomal aberrations. The pink, green and yellow colors indicate the raw signal for each individual chromosome probe, while the blue signal represents the normalized probe signal which is used to identify copy number and aberrations (if any). (**D**) Western blots demonstrating SynGAP protein expression from iNeuron or iPSC homogenate. Total refers to signal from an antibody that detects all splice variants and α2 refers to signal from an antibody that detects only a specific C-terminal splice variant. (**E**) Quantification of relative intensity of bands normalized to total protein signal. One-way ANOVA with a Kruskal-Wallis test multiple comparisons test H(3)=12.29, p=0.0001; WT#6 vs KO#4: p=0.1876; WT#6 vs WT#30: p>0.9999; WT#6 vs KO#38: p=0.0140; KO#4 vs WT#30: p=0.5258; KO#4 vs KO#38: p>0.9999; WT#30 vs KO#38: p=0.0561. n=4 per group. In box-and-whisker plot, the center, boxes and whiskers represent the median, interquartile range, and min to max, respectively. **(F)** Indels from each clone identified from whole exome sequencing analysis. Indels were identified by clonal sequence differences from the original Cas9 hiPSCs (reference sequence). Indel threshold was determined by at least 50% of the reads differing from the reference sequence with a minimum of at least ten reads. Indels w/ frequency above 0.8 were used to determine frequency of homozygous varaints. (**G**) Normalized mapped reads from the entire coding sequence of the *SYNGAP1* gene in the four clones hiPSCs. Red arrow denotes predicted Cas9 cut site. Numbers reflect clonal reads relative to Cas9 hiPSC reads. (**H**) Normalized mapped reads for the same samples around the Cas9 target sequence. (**I**) Genomic PCR to amplify DNA sequence flanking the Cas9 target site.

We next performed whole exome sequencing (WES) to quantify the genetic differences among the clones. In general, the four clones had very little genetic drift across the protein coding portion of the genome. We observed only a few high confidence exonic small indels in each of the four CRISPR clones **(Table 1)**. None of these indels were shared within the same gene and none of them appeared to be homozygous, except for the two unique indels identified in Exon7 of *SYNAGP1* **(Fig. 1F)**. Thus, unbiased read-mapping of WES identified the sequences used to select the two “KO” clones and these sequences appeared to be the most significant deviations amongst the four clones (**Fig. 1A-B; Table 1)**. Therefore, these four clones are essentially isogenic, with the exception of the homozygous disruptive variants present in *SYNGAP1*. We next performed in-depth mapping of *SYNGAP1* exons to further characterize potential off-target effects of Cas9 genome editing **(Fig. 1G)**. Comparing normalized reads of each of the four clones relative to the original iPSC line revealed that *SYNGAP1* exon structure was largely intact. However, we did observe a ~50% reduction in mapped reads in the targeted exon of KO clone #38. Flanking exons had normal read depths **(Fig. 1H)**. This was suggestive of a large deletion that encompassed exon 6/7, but was less than 4Kb in size. Genomic PCR failed to detect a band shift **(Fig. 1I)**, though PCR amplification in this region was limited to ~2.2Kb. Thus, for clone #38, there were likely two distinct Indels in each of the *SYNGAP1* copies. Copy 1 contained an 8bp deletion **(Fig. 1B; Table 1)**, while the other copy likely contained an undefined indel (<4Kb) that prevented amplification by traditional PCR. In contrast, clone #4 appeared to contain a bi-allelic single base deletion in exon 7. In each case, these indels produced nominal SynGAP protein in induced neurons **(Fig. 1D-E)**. We conclude that the two “KO” clones, when paired with the isogenic “WT” clones, were suitable to determine the impact of SynGAP loss-of-function on human neuron development and function.

**Table 1.**
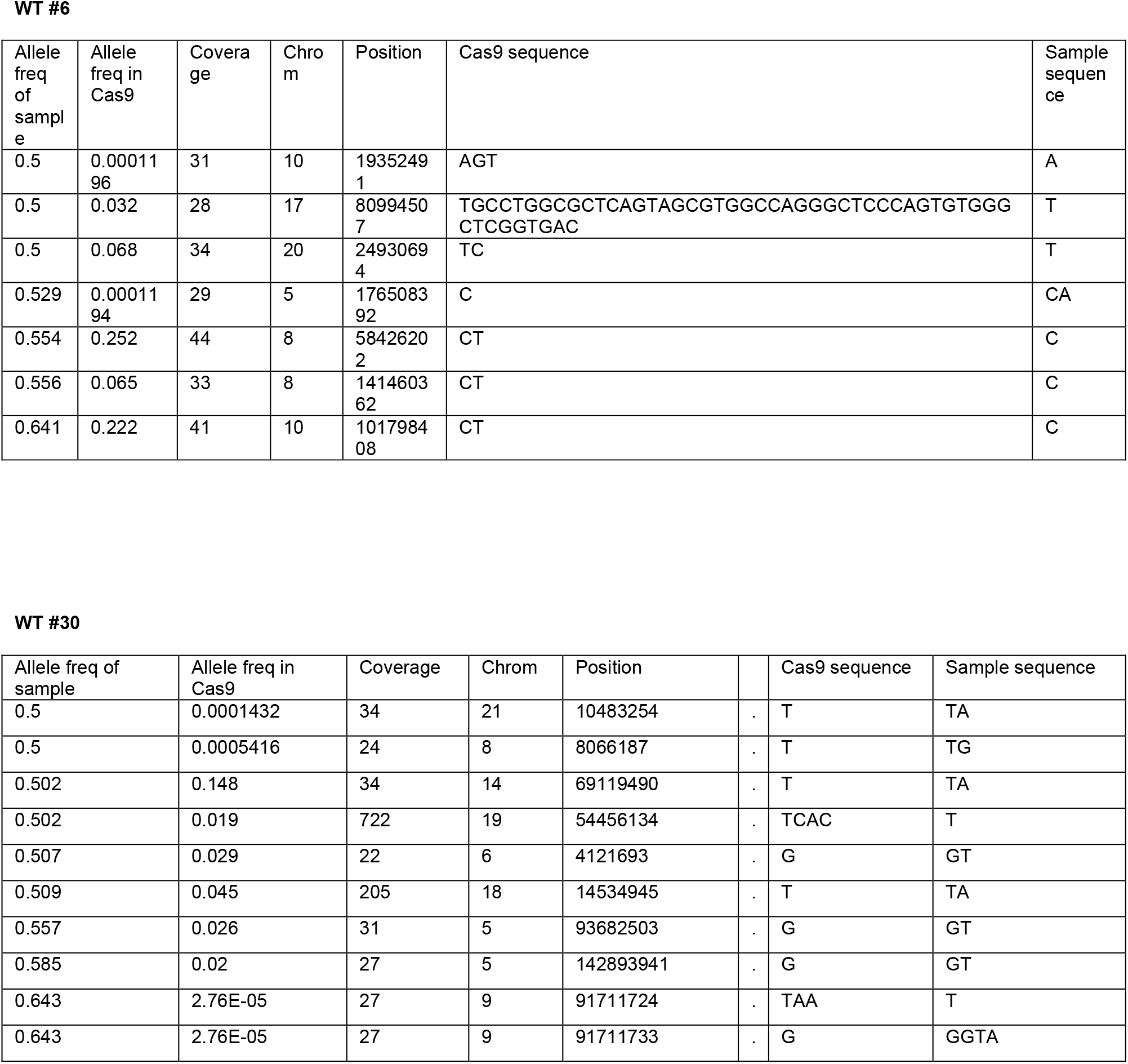
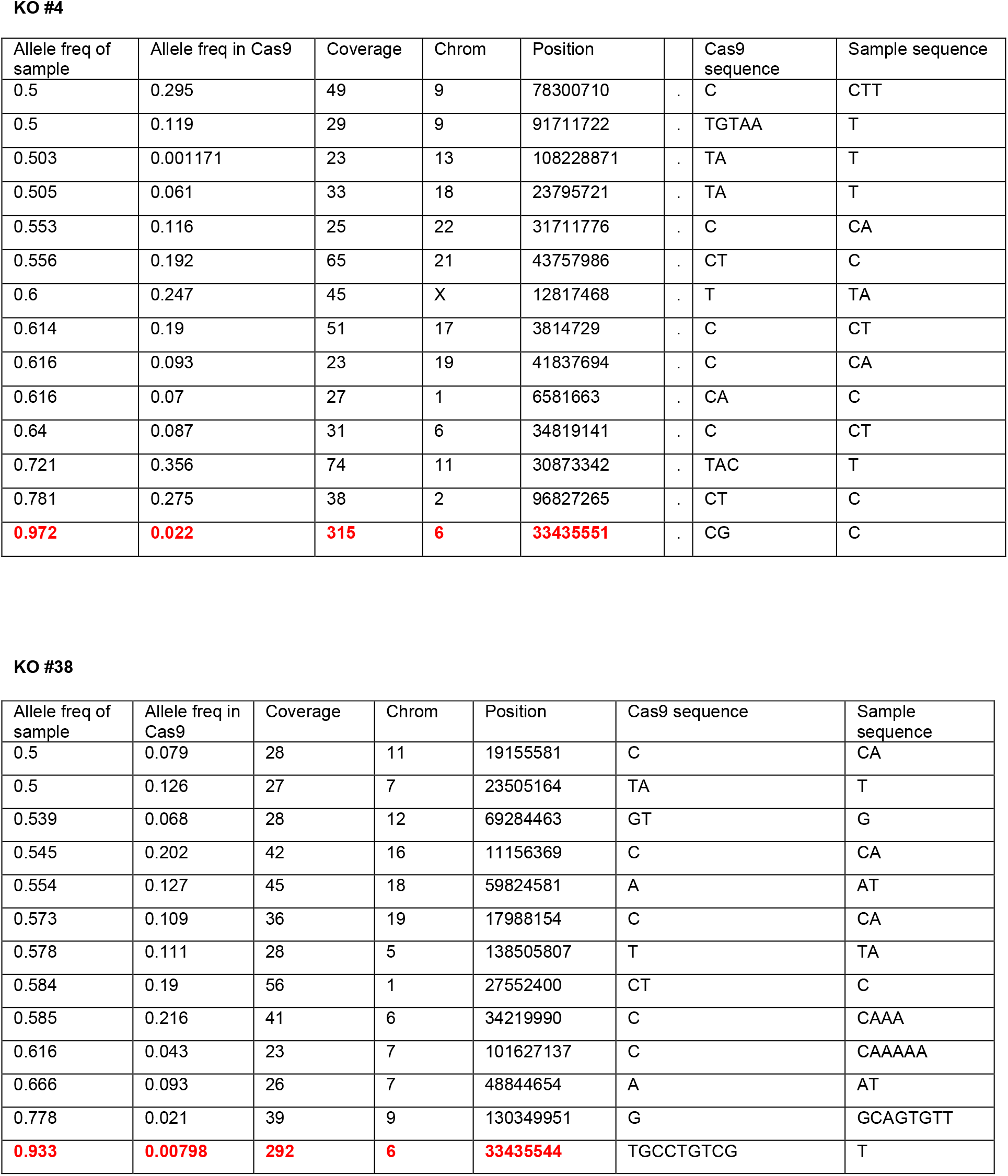
Indels present in the clonal SYNGAP1 iPSC lines (relative to starting material)

*Syngap1* loss-of-function in rodent neurons disrupts the maturation rate of dendrites and synapses. Therefore, we examined dendritic morphogenesis in developing iNeurons produced from each of the four human iPSC clones. Dendritic morphology was measured at day *in vitro* (DIV) 45 by tracing dendrites of sparsely labeled eGFP-positive iNeurons (**Fig. 2A**). Relative to each isogenic control line, total dendritic fields were substantially larger in iNeurons derived from *SYNGAP1*-KO clones. This difference was observed at the level of individual clones (**Fig. 2A-B**) and when clones were grouped by genotype (**Fig. 2B**). Examination of the length by dendritic category (e.g. primary) revealed that, compared to WT clones, KO clones generally had longer primary and secondary dendrites (**Fig. 2A, C-E)**. The lack of a clonal difference within tertiary dendrites likely reflected a lower statistical power, as many neurons lacked these structures. In contrast to length, the complexity of dendritic arbors was unaffected by *SYNGAP1* disruption. Clonal and genotype effects of *SYNGAP1* were not observed for total dendrites (**Fig. 2F**). Moreover, no *SYNGAP1* effects were observed for each dendrite subtype (**Fig. 2G-I).**

**Figure 2.**
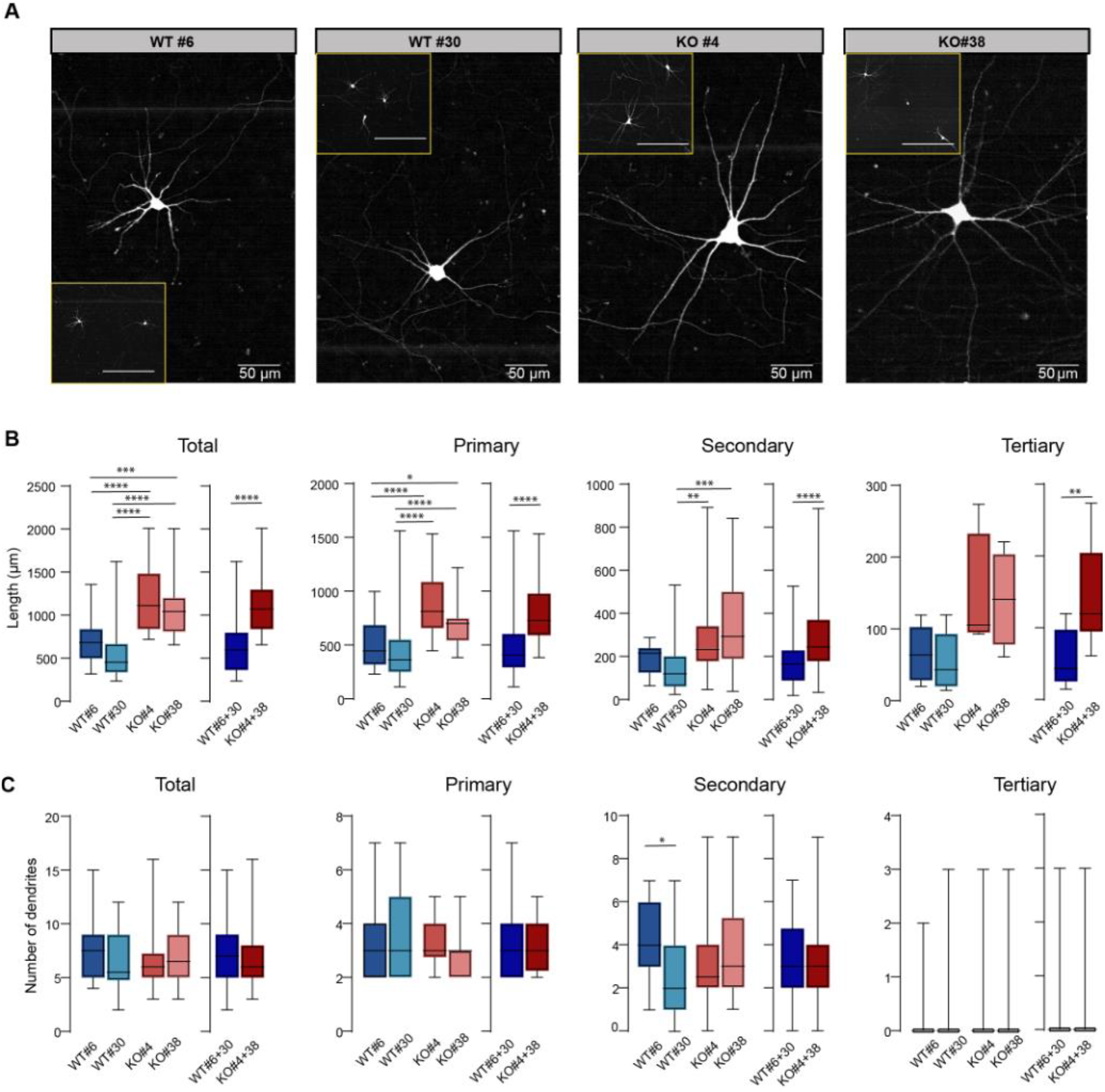
Increased dendrite length in iNeurons derived from KO iPSC clones. (**A**) presentative images of eGFP-expressing iNeurons from the four different clones at DIV45 et images scale bars: 200 μm). (B-E) Histograms depicting average length per cell of total, primary (**C**), secondary (**D**) and tertiary dendrites (**E**) of the four clones (*Total dendrites* - nal analysis H=54.81, p<0.0001; N=30 cells per clone; Genotype analysis U=436, p<0.0001; 60 cells per genotype; *Primary dendrites* – Clonal analysis H=49.71, p<0.0001, N=30 cells clone; Genotype analysis U=545, p<0.0001, N=60 cells per genotype; *Secondary dendrites* lonal analysis H=20.45, p<0.0001; N=30 cells for WT#6 and KO#38; N=26 cells for WT#30; 27 cells from KO#4; Genotype analysis U=880, p<0.0001; N=56 cells from WT genotype; N=57 cells from KO genotype; *Tertiary dendrites* – Clonal Analysis, H=7.115, p=0.0683; N=6 dendrites of 30 cells from WT#6, n=5 dendrites of 30 cells from WT#30, n=4 dendrites of 30 cells from KO#4 and n=4 dendrites of 30 cells from KO#38; Genotype analysis, U=73.55, p=0.0068; N=11 cells from WT genotype; N=8 cells from KO genotype). (F-I) Graphs showing average number of dendrites per cell of total (**F**), primary (**G**), secondary (**H**) and tertiary (**I**) dendrites of the four clones (*Total dendrites* – Clonal analysis, H=5.957, p=0.1137; N=30 cells per clone; Genotype analysis, U=1613, p=0.3222; N=60 cells per genotype; *Primary dendrites* – H=1.680, p=0.6413, n=30 cells per clone; Genotype analysis, U=1639, p=0.3755, n=60 cells per genotype; *Secondary dendrites –* Clonal analysis, H=10.72, p=0.0133, n=30 cells per clone for clone comparisons; Genotype analysis, U=1689, p=0.5552, n=60 cells per genotype; *Tertiary dendrites –* Clonal analysis, H=0.4531, p=0.9291, n=30; Genotype analysis, U=1731, p=0.6129, n=60 cells per genotype). In box-and-whisker plots, the center, boxes and whiskers represent the median, interquartile range, and min to max, respectively. *p<0.05, **p<0.01,****p<0.0001.

Tracing studies suggested that reduced SynGAP expression leads to iNeurons with larger dendritic fields. To confirm this, we performed an orthogonal analysis, consisting of immunocytochemical labeling of dendritic and synaptic proteins, in neurons derived from one pair of isogenic WT or KO iPSCs **(Fig. 3A-B)**. The MAP2 area was enhanced in KO cultures **(Fig. 3A-C)** and was not due to more KO neurons plated in these cultures **(Fig. 3A-E)**. Cultured neurons with longer dendrites would be expected to have an increase in absolute numbers of postsynaptic structures. Indeed, absolute numbers of PSD95 and GLUA1 structures were also increased in the KO culture **(Fig. 3A-C)**. The effect of genotype on synaptic labeling was still significant, albeit with a much smaller effect size, when PSD95 and GLUA1 structures were normalized to MAP2 area **(Fig. 3F)**. These labeling studies support the idea that disrupting SynGAP expression results in cultures comprised of larger neurons with more postsynaptic structures.

**Figure 3.**
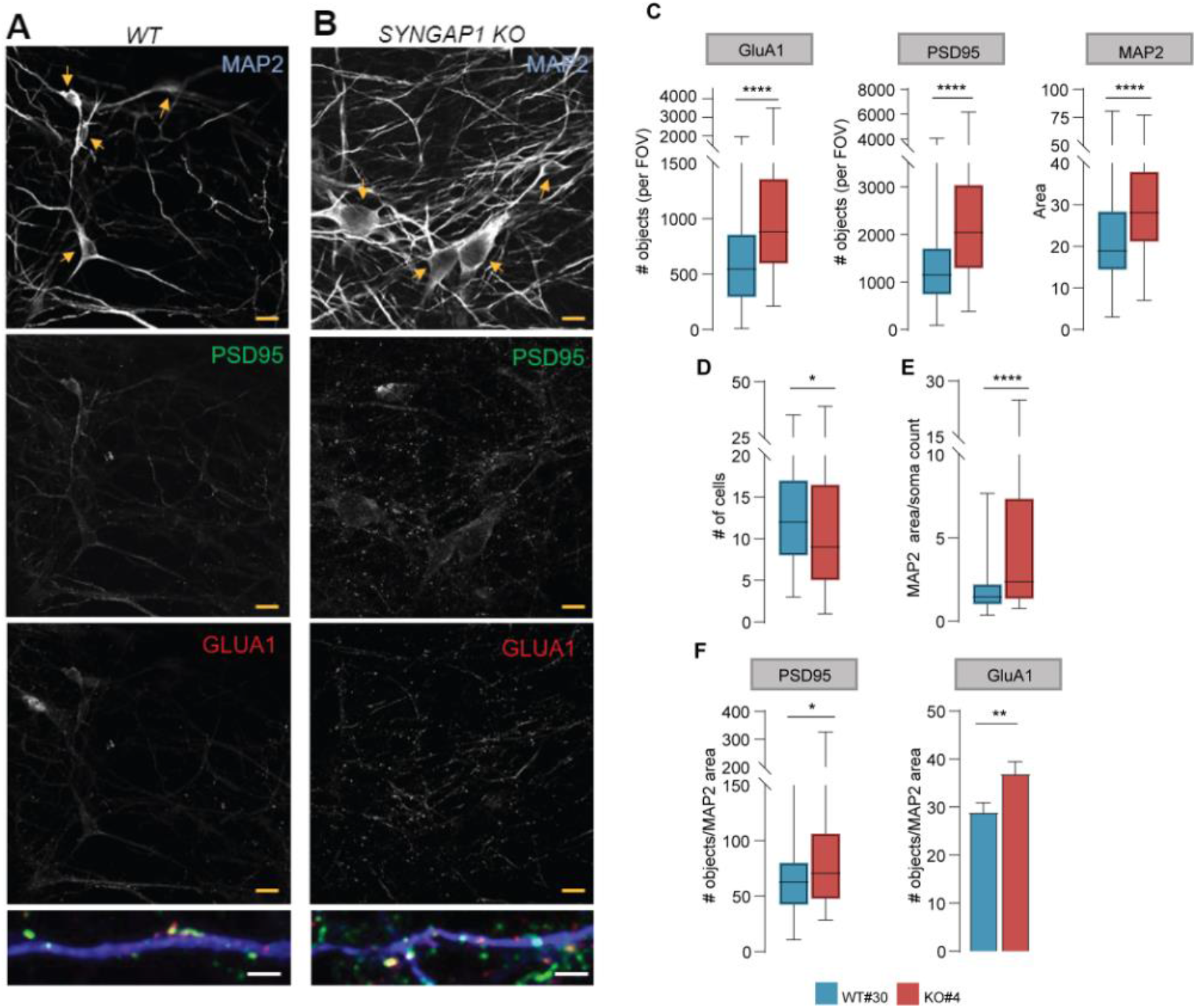
Increased dendritic area and more numerous postsynaptic structures in *SYNGAP1* KO iNeurons. (A-B) Representative images showing MAP2 labeling (top), PSD95 labeling (middle), and GluA1 labeling (middle-bottom), and merge of MAP2, PSD95 and GLUA1 (bottom) of iNeurons from WT#30 (**A**) and KO#4 (**B**) at DIV45. Yellow arrows indicate cell bodies. Yellow scale bar is 10 μm and white is 2 μm. (**C**) Graphs showing MAP2 area and number of PSD95 objects (punctate labeling) per field of view (FOV) in WT#30 and KO#4. (MAP2 comparison: U=2181, p<0.0001 N=85 images for WT#30 and 81 images for KO#4; PSD95 comparison: U=1891, p<0.0001 N=85 for WT#30 and 81 for KO#4; GluA1 comparison: U=1999, p<0.0001 N=85 for WT#30 and 81 for KO#4) (**D**) Averaged number of cells in FOVs WT#30 and KO#4 (U=2746, p=0.0240 N=85 for WT#30 and 81 for KO#4). (**E**) MAP2 area by soma count (U=2097, p<0.0001 N=85 for WT#30 and 81 for KO#4). (**F**) Graphs showing quantification of PSD95 and GluA1 expression in WT#30 and KO#4 normalized to MAP2 area (PSD95 objects/MAP2 area comparison: U=2752, p=0.0255 N=85 for WT#30 and 81 for KO#4; GluA1 objects/MAP2 area comparison: t_(164)_=2666, p=0.0084 N=85 for WT#30 and 81 for KO#4). In box-and-whisker plots, the center, boxes and whiskers represent the median, interquartile range, and min to max, respectively. Bar graph represents mean ± SEM. *p<0.05, **p<0.01 and ****p<0.0001

The observation of larger iNeurons with increased numbers of postsynaptic structures prompted us to investigate the functional maturation of iNeurons with reduced SynGAP protein expression. Intrinsic membrane properties and the onset of glutamatergic synaptic activity are two measures that are developmentally regulated in Ngn2-induced neurons (Zhang et al., 2013). To test the idea that reducing SynGAP expression alters the maturation of iNeurons, we performed whole-cell voltage-clamp recordings at two developmental time points (*DIV20-30 and DIV40-50*; **Fig. 4A-B**). At DIV20-30, intrinsic membrane properties of all clones were characteristic of immature neurons (i.e. relatively low capacitance and high input resistance; **Fig. 4C-D**). We did not observe clonal or genotype differences in resting membrane potential, capacitance, or resistance at this time point (**Fig. 4C-E**). However, we did observe that neurons made from *SYNGAP1*-KO hiPSCs showed earlier synaptic activity during development. Although some iNeurons from all clones exhibited miniature excitatory postsynaptic currents (*m*EPSCs) at this time point (**Fig. 4F**), the proportion of *m*EPSC-expressing iNeurons was significantly increased in KO clones (**Fig. 4G**). When grouping iNeurons by genotype, KO neurons were almost twice as likely to express miniature events (**Fig. 4G**). *m*EPSC frequency was low and variable at this early time point, making it difficult to compare clones or even genotypes **(Fig. 4H-I)**. In contrast, *m*EPSC amplitude was less variable. There appeared to be a weak clonal and genotype effect on *m*EPSC amplitude. Both KO amplitude populations exhibited a rightward shift compared to the two WT populations **(Fig. 4J)**. When clonal data was collapsed by genotype, a robust statistical effect emerged at the level of individual events and at the level of cellular population means **(Fig. 4J).**

**Figure 4.**
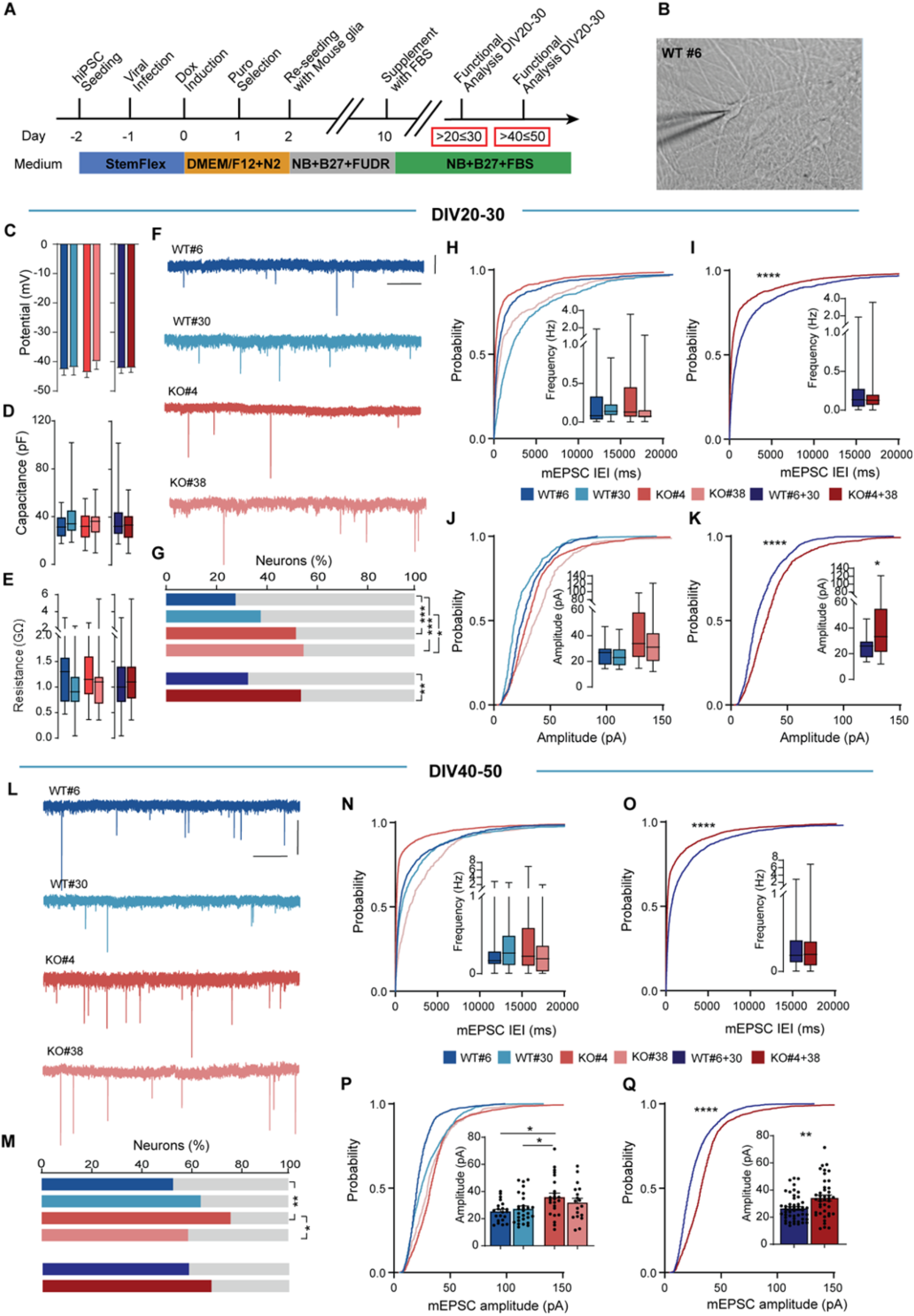
*SYNGAP1* expression in human iNeurons regulates excitatory synapse function. (**A**) Flow diagram of iNeuron generation from WT and *SYNGAP1* KO iPSCs for whole-cell electrophysiological experiments (recording days within red boxes). (**B**) Representative DIC image of patched iNeurons cells from WT#6. (C-E) Bar graphs representing intrinsic membrane properties measured at DIV20-30 as resting membrane potential (**C**), capacitance (**D**) and input resistance (**E**) from the four clones (*Membrane potential* – Clonal analysis, F_(3.95)_=0.5132, p=0.6742, n=29 cells from WT#6, 31 cells from WT#30, 34 cells from KO#4 and 21 cells from KO#38;Genotype analysis, t_(97)_=0.08684, p=0.9310, n=51 cells for WT#6+30 and 48 for KO#4+38; *Capacitance* – Clonal analysis, H=3.123, p=0.3730, n=29 cells from WT#6, 31 cells from WT#30, 34 cells from KO#4 and 21 cells from KO#38; Genotype analysis, U=1584, p=0.5093; N=62 cells from WT#6+30 and 55 cells from KO#4+38; *Membrane resistance*- Clonal analysis, H=4.259, p=0.2348, n=28 cells from WT#6, 31 cells from WT#30, 34 cells from KO#4 and 21 cells from KO#38; Genotype analysis, U=1546, p=0.5619; N=60 cells from WT#6+30 and 55 cells from KO#4+38). (**F**) Representative traces of mEPSCs of iNeurons from WT and KO clones at DIV20-30. Scale bars 2 s, 20pA. (**G**) Percentage of successful observations of mEPSCs in iNeurons from the four clones at DIV20-30 (Clonal analysis, p=0.0008 for KO#4 vs WT#6; p=0.0002 for KO#38 vs WT#6, p=0.0644 for KO#4 vs WT#30, n=20 for KO#4 and 18 for WT#30, resp; p=0.0231 for KO#38 vs WT#30; N= 12 cells from WT#6, 18 cells from WT#30, 20 cells from KO#4 and 19 cells from KO#38; Genotype analysis, p=0.0042 for KO#4+38 vs WT#6+30, n=30cells from WT#6+30 and 39 cells from KO#4+38). (H- I) Cumulative plots of mEPSC interevent-interval and frequency (inset) of the different clones individually (**H**) and grouped by genotype (**I**) at DIV20-30 (Clonal analysis, H=1.910, p=0.5912, n=12 cells from WT#6, 18 cells from WT#30, 20 cells from KO#4 and 19 cells from KO#38; Genotype analysis, U=504.5, p=0.5607; N=29 cells from WT#6+30 and 38 cells from KO#4+38, K-S test D=0.2660, p<0.0001, n= 951 events from 29 cells from WT#6+30, n= 1559 events from 38 cells from KO#4+38). (J-K) Cumulative probability plots of mEPSC amplitude of the different clones individually (**J**) and grouped by genotype (**K**) at DIV20-30 (Clonal analysis, H=7.565, p=0.0559, n=15 cells from WT#6, 20 cells from WT#30, 19 cells from KO#4 and 19 cells from KO#38; Genotype analysis, U=504.5, p=0.5607; N=29 cells from WT#6+30 and 38 cells from KO#4+38, K-S test D=0.2660, p<0.0001, n= 981 events from 35 cells from WT#6+30, n= 1601 events from 38 cells from KO#4+38). (**L**) Representative traces of mEPSCs of iNeurons from WT and KO clones at DIV40-50. Scale bars 2 s, 20pA. (**M**) Percentage of successful observations of mEPSCs in iNeurons from the four clones at DIV40-50 (Clonal analysis, p=0.0011 for WT#6 vs KO#4; p=0.4764 for WT#6 vs KO#38; p=0.0892 for WT#30 vs KO#4; p=0.5612 for WT#30 vs KO#38; p=0.1511 for WT#6 vs WT#30; p= 0.0154 for KO#4 vs KO#38; N= 21 cells from WT#6, 29 cells from WT#30, 25 cells from KO#4 and 18 cells from KO#38; Genotype analysis, p=0.2399 for WT#6+30 vs KO#4+38; N=50 cells from WT#6+30 and 43 cells from KO#4+38; Effect of time, p=0.0004 for WT#6+30 p40 vs p20; p= p=0.0592 for KO#4+38 p40-50 vs p20-30. (N-O) Cumulative probability plots of mEPSC interevent-interval (IEI) and frequency (inset) of the different clones individually (**N**) and grouped by genotype (**O**) at DIV40-50 (Clonal analysis, H=2.874, p=0.4115, n=21 cells from WT#6, 28 cells from WT#30, 24 cells from KO#4 and 18 cells from KO#38; Genotype analysis, U=970.5, p=0.644; N=49 cells from WT#6+30 and n=42 cells from KO#4+38, K-S test D=0.2763, p<0.0001, n= 2182 events from 49 cells from WT#6+30, n= 2498 events from 42 cells from KO#4+38). (P-Q) Cumulative probability plots of mEPSC amplitude of the different clones individually (**P**) and grouped by genotype (**Q**) at DIV40-50 (Clonal analysis, F_(3.87)_=3.73, p=0.0142, p=0.0187 for KO#4 vs WT#6, p=0.0499 for KO#4 vs WT#30, p=0.9407 for WT#6 vs WT#30; p=0.3151 for WT#6 vs KO#38; p=0.5696 for WT#30 vs KO#4 and p=0.70 for KO#4 vs KO#38; Genotype analysis, t_(89)_=3.121, p=0.0024, n=49 cells for WT#6+30 and 42 for KO#4+38, K-S test D=0.2990, p<0.0001, n= 2254 events from 49 cells from WT#6+30, n= 2554 events from 42 cells from KO#4+38). In box-and-whisker plots, the center, boxes and whiskers represent the median, interquartile range, and min to max, respectively. Bar graphs represent mean ± SEM. *p<0.05, **p<0.01

We next analyzed synaptic activity in more mature iNeurons (*DIV40-50*; **Fig. 4L)**. As a population, neurons derived from WT clones were roughly twice as likely to express synaptic activity at this time point compared to younger neurons of the same genotype, indicative of substantial neuronal maturation during this period **(Fig. 4G, M)**. However, this effect was less pronounced in KO neurons **(Fig. 4G, M)**. There was a significant effect of time on the proportion of neurons expressing synaptic activity in WT neurons, but this effect was absent in KO iNeurons (**Fig. 4M)**. There was no longer an effect of genotype on the proportion of neurons with synaptic activity at the more mature stage of development. Within the population of neurons with synaptic events, we measured mEPSC frequency and amplitude. The frequency of events was highly variable in these populations **(Fig. 4N-O)**, which made it difficult to draw clear conclusions across clones and genotypes. There was a trend toward more frequent events in combined KO populations, though these trends were not apparent when looking at individual clones. With respect to amplitude **(Fig. 4P-Q)**, we once again observed a weak effect of clone and genotype at this timepoint that was consistent with observations from developmentally younger iNeurons. Neurons from both KO clones appeared to have slightly larger events compared to those from WT iNeurons. This effect was apparent in comparisons of mEPSC distributions of all events **(Fig. 4P-Q),** and in the much less sensitive approach of comparing event means from individual neurons **(Fig. 4P-Q,** *inset*).

The effect of SynGAP expression on iNeuron *m*EPSC frequency and amplitude was somewhat consistent across developmental time points, but the effect sizes, when present, were relatively small. To determine if these effects were reproducible, we performed an additional experiment on iNeurons produced from the same clones. Data for this experiment was collected from a completely new hiPSC expansion and neuronal induction procedure. In this additional experiment, we observed similar effects of SynGAP expression on intrinsic membrane properties and *m*EPSCs **(Fig. 5A-G)**. *SYNGAP1* deletion did not affect the resting membrane potential, input resistance, or capacitance at the clonal or genotype level **(Fig. 5A-C)**. Analysis of mEPSC frequency from each of the clones revealed a trend for increased frequency from neurons with disruptive *SYNGAP1* variants **(Fig. 5D)**. The two KO clones have a greater frequency of mEPSCs when looking at cumulative probability distributions and this drove an effect at the genotype level **(Fig. 5E)**. A statistical effect was not present when comparing cellular means of mEPSC frequency. For mEPSC amplitude, the clonal and genotype effects were clearer compared to frequency measures. The cumulative distribution for mEPSC amplitudes for all events clearly shifted to larger values in both KO clones **(Fig. 5F)**. This drove a substantial and highly significant shift in the disruption at the genotype level **(Fig. 5G)**. We did not observe an effect on population means when looking at cellular averages. However, the power for this experiment was lower than the one presented in Figure 4. Taken together, we conclude that reducing SynGAP expression in Ngn2 iNeurons leads to weak, but reproducible, effects on mEPSC amplitude. Effects on frequency were unclear due to high variability.

**Figure 5.**
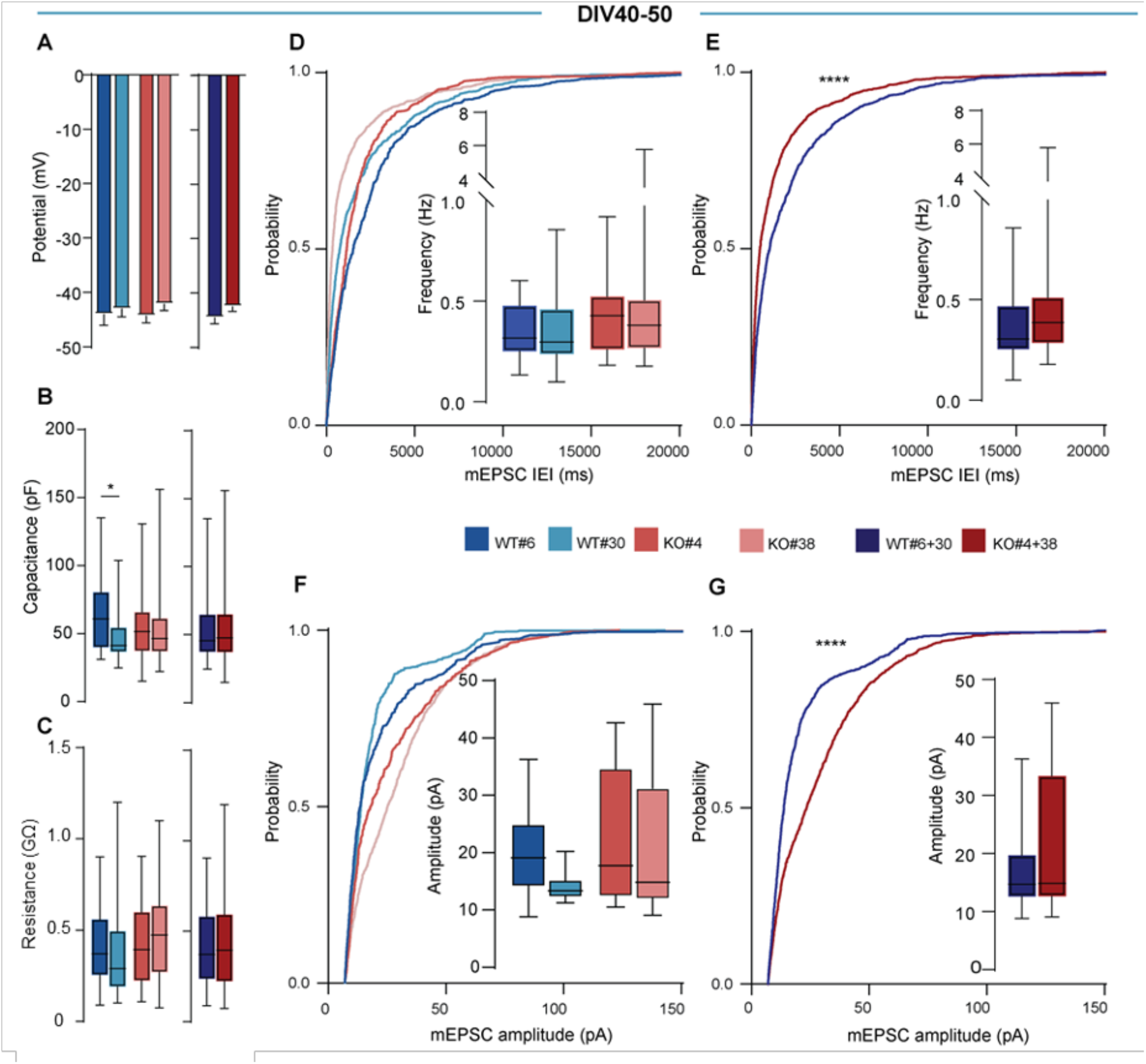
Reproducibility of *SYNGAP1*-mediated effects on iNeuron excitatory synapse function. (A-C) Graphs showing resting membrane potential (**A**), capacitance (**B**) and input resistance (**C**) from the four clones at DIV40-50 (*Membrane potential* – Clonal analysis, F_(3.54)_=0.5456, p=0.6532, n=11 cells from WT#6, 16 cells from WT#30, 13 cells from KO#4 and 18 cells from KO#38; Genotype analysis, t_(56)_=1.215, p=0.2295, n=27 cells for WT#6+30 and 21 for KO#4+38; *Capacitance* – Clonal analysis, H=9.091, p=0.0281, p=0.0318 for WT#6 vs WT#30, n=28 cells from WT#6, 41 cells from WT#30, 34 cells from KO#4 and 50 cells from KO#38; Genotype analysis, U=2828, p=0.7973; N=69 cells from WT#6+30 and 84 cells from KO#4+38; *Membrane resistance*- Clonal analysis, H=4.738, p=0.1920, n=28 cells from WT#6, 41 cells from WT#30, 34 cells from KO#4 and 50 cells from KO#38; Genotype analysis, U=2896, p=0.9949; N=69 cells from WT#6+30 and 84 cells from KO#4+38).(D-E) Cumulative plots of mEPSC interevent-interval (IEI) and frequency (inset) of the different clones individually (**D**) and grouped by genotype (**E**) at DIV40-50 (Clonal analysis, H=1.663, p=0.6452, n=11 cells from WT#6, 9 cells from WT#30, 8 cells from KO#4 and 13 cells from KO#38; Genotype analysis, U=164, p=0.2382; N=20 cells from WT#6+30 and n=21 cells from KO#4+38, K-S test D=0.1744, p<0.0001, n= 1199 events from 20 cells from WT#6+30, n= 1512 events from 21 cells from KO#4+38). (F-G) Cumulative probability plots of mEPSC amplitude of the different clones individually (**F**) and grouped by genotype (**G**) at DIV40-50 (Clonal analysis, H=3.080, p=0.3795, n=11 cells from WT#6, 9 cells from WT#30, 8 cells from KO#4 and 13 cells from KO#38; Genotype analysis, U=182, p=0.4773; N=20 cells from WT#6+30 and n=21 cells from KO#4+38, K-S test D=0.2954, p<0.0001, n= 1085 events from 20 cells from WT#6+30, n= 1396 events from 21 cells from KO#4+38). In box-and-whisker plots, the center, boxes and whiskers represent the median, interquartile range, and min to max, respectively. Bar graphs represent mean ± SEM. *p<0.05.

Our data demonstrate that reducing SynGAP expression results in larger iNeurons that exhibit early synaptic maturity. Therefore, we hypothesized that reducing SynGAP expression would also influence the development of network activity in cultured iNeurons. To test this, we measured spontaneous distributed network activity in cultures derived from KO and WT clones using a multielectrode array (MEA) system (**Fig. 6A-C**). Recordings of the same cultures were performed over the course of several weeks, which enabled *in vitro* measurements of network spiking activity during neuronal development. From as early as week 2, we observed evidence of spiking activity in cultures derived from each of the iPSC clones. However, both *SYNGAP1* KO clones exhibited substantially increased firing rates compared to isogenic controls. The enhanced firing rate in KO iNeurons emerged progressively and was sustained through week six in culture at both clonal (**Fig. 6D**) and genotype levels (**Fig. 6E**). Next we measured bursting activity in each of the four clones. We observed significantly elevated neuronal bursts in KO vs control neurons **(Fig. 6F-G)**. Quantification of distributed network connectivity demonstrated that KO neuronal cultures displayed different degrees of neural network activity, observed as “network bursts”, as early as 3 weeks of maturation. Enhanced network bursting activity in KO cultures relative to WT controls was observed at both the clonal (**Fig. 6H**) and genotype levels (**Fig. 6I**). Thus, *SYNGAP1* expression substantially influences the dynamics of cellular activity in developing neuronal networks.

**Figure 6.**
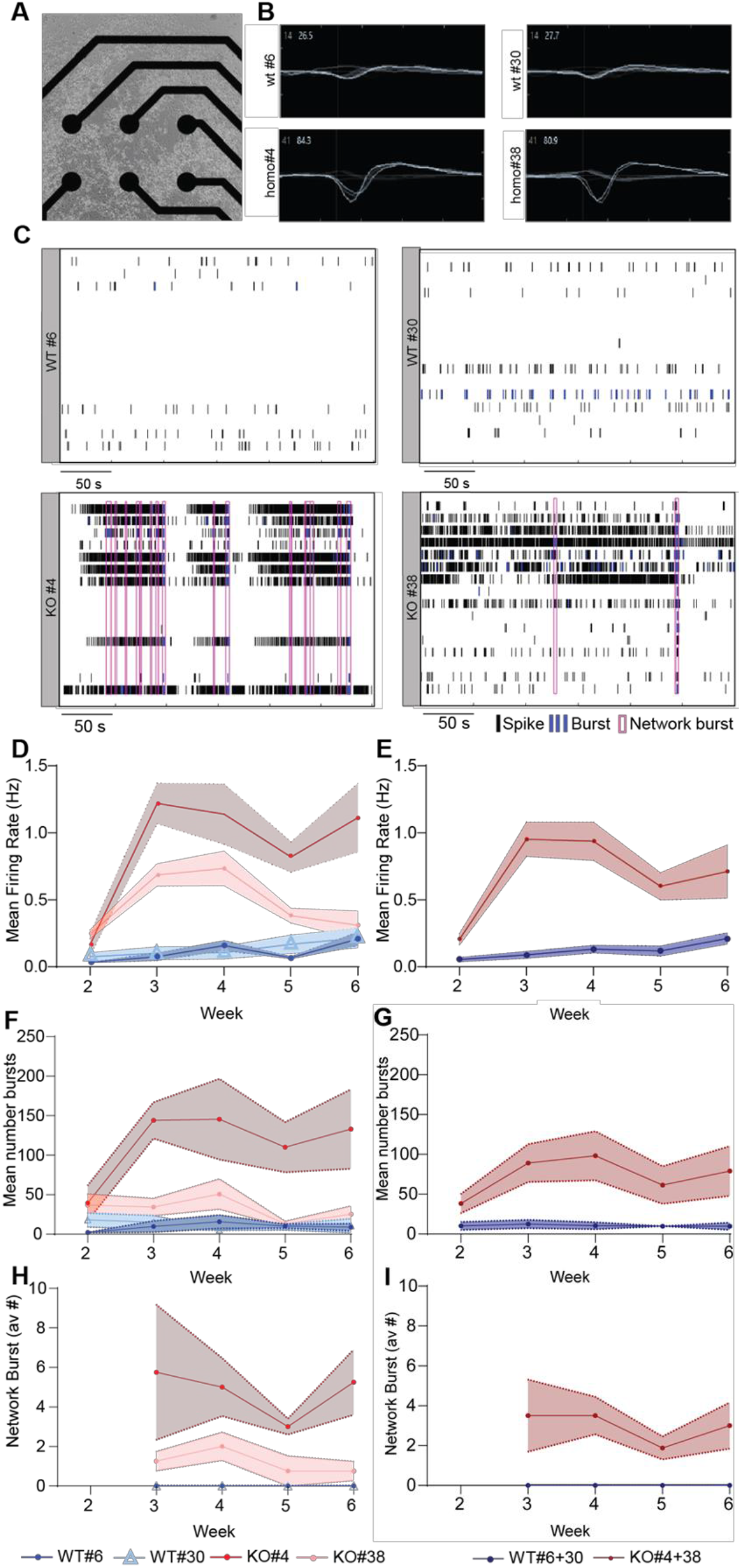
Earlier onset and elevated levels of network activity in *SYNGAP1* KO iNeurons. (**A**) Representative bright-field image of 1-week old iNeurons differentiated from iPSC-derived NPCs plated on a 16-electrode array of an MEA well. Spontaneous action potentials were recorded from the homozygous *SYNGAP1* null (Homo#4 and #30) and control (WT#6 and #30) neurons. (**B**) Representative wave forms of spiking behavior from a single electrode for each Homo and WT neuronal culture. (**C**) Representative temporal raster plots of KO iNeurons (KO#4 and #38) and WT isogenic control iNeurons (WT#6 and #30) over 5-minuntes of continuous recording during culture week 3. (D-E) Cumulative plots of mean firing rates for all four clones individually (**D**) and grouped together by genotype (**E**), along a developmental timeline. (F-G) Cumulative plots of average number of bursts for individual clones **(F)** and grouped together by genotype **(G)**. (H-I) Cumulative plots of average number of network bursts for all clone individually **(H)** and grouped together by genotype **(I)**. KO neurons display synaptic connections as early as week 3 of maturation compared to the WT controls. (*Week 2* – Clonal analysis, H=9.331, p=0.0096, post-hoc comparisons p=0.0227 for KO#38 vs WT#6, p>0.999 for WT#6 vs WT#30; p=0.2697 for WT#6 vs KO#4; p>0.9999 for KO#4 vs KO#38 and p=0.999 for WT#30 vs KO#4 and p=0.3803 for WT#30 vs KO#38; Genotype analysis, U=6, p=0.0047; *Week 3* – H=12.73 p<0.0001, post-hoc comparisons p=0.0140 for KO#4 vs WT#6, p=0.0227 for KO#4 vs WT#30;Genotype analysis U=0, p=0.0002; *Week 4* – Clonal analysis, H=12.29 p=0.0001, post-hoc comparisons p=0.01410 for KO#4 vs WT#30; Genotype analysis, U=0, p=0.0002; *Week 5* – Clonal analysis, H=11.89 p=0.0004, post-hoc comparisons p=0.0084 for KO#4 vs WT#6; Genotype analysis, U=2, p=0.0006; *Week 6* – H=9.088, p=0.0117; post-hoc comparisons p=0.0451 for KO#4 vs WT#6; Genotype analysis, U=10, p=0.0207). N= 4 replicas for WT#6, WT#30, KO#4 and KO#38; N=8 replicas from WT#6+30 and KO#4+38. For each clone, four replicates of iNeurons were plated and differentiated concurrently. Bar graph represents mean ± SEM. *p<0.05, **p<0.01, ***p<0.001 and ****p<0.0001.

## Discussion

We produced iNeurons from human hiPSCs with a disrupted *SYNGAP1* gene in an effort to understand how this gene shapes human neuron development and function. This is an important research question given that pathogenic *SYNGAP1* variants cause a complex neurodevelopmental disorder defined by early-onset epilepsy, cognitive impairment, and autistic features (Hamdan et al., 2011; Jimenez-Gomez et al., 2019; Vlaskamp et al., 2019; Satterstrom et al., 2020). We found that *SYNGAP1* regulates the postmitotic maturation of dendrites and synapses from human iNeurons. Cas9-mediated disruption of *SYNGAP1* expression enhanced dendritic morphogenesis, accelerated the acquisition of synaptic activity, and drove increased spiking activity measured in functionally connected two-dimensional iNeuron cultures. Our data indicate that loss of SynGAP protein expression was responsible for the dendrite and synapse maturation phenotypes observed in these cultures. Indeed, we observed consistent structural phenotypes at the level of individual clones that were subsequently grouped by genotype. Whole exome sequencing demonstrated that the only shared variants between the two KO clones were indels in the *SYNGAP1* gene, and immunoblotting confirmed that iNeurons derived from KO clones expressed nominal levels of SynGAP protein. Altered dendritic maturation was supported by data obtained from orthogonal experimental measures. We observed longer dendrites in eGFP-positive iNeurons, and an increased dendritic area measured from endogenous MAP2 signal. iNeurons derived from the KO hiPSC clone also exhibited an increase in the absolute density of postsynaptic structures, a finding consistent with a neuronal culture populated with neurons containing longer dendrites. Given that the length of dendrites and the density of postsynaptic structures in iNeurons increases over time in culture (Zhang et al., 2013), these data support the conclusion that SynGAP expression regulates the maturation rate of dendritic and synaptic structures in human iNeurons. This conclusion was also supported by clonal and genotype differences in synaptic activity between WT and KO iNeurons. Individual iNeurons have been shown to gradually acquire synaptic activity in the first several weeks in culture (Zhang et al., 2013; Nehme et al., 2018). However, we found that KO neurons expressed synaptic activity earlier in development compared to WT neurons.

Distributed neuronal activity, measured by MEA analysis, confirmed that structural maturation of dendrites and early functional expression of synapse activity translated into increased network activity in KO cultures. Similar to what we observed in dendrites and synapses, measures of network activity normally observed in more mature WT cultures appeared at much earlier stages of development in neurons developed from KO clones. Activity was already substantially greater in cultures derived from KO clones at two weeks, a time in development when there is very little activity present in WT cultures. In addition, statistical analysis of network activity that considered time as a factor demonstrated that the trajectory of neuronal activity was distinct in KO cultures compared to WT controls (Fig. 6). Indeed, activity increased at a much greater rate in KO cultures, compared to WTs, over the first several weeks of development. Networks formed from iNeurons exhibited bursting behavior as a function of time *in vitro*, with older cultures exhibiting more robust bursting behavior (Fischer). Network bursting is driven in part by increased functional synaptic connectivity among neurons (Suresh et al., 2016; Nehme et al., 2018). KO neurons extended dendrites more quickly and had greater numbers of postsynaptic structures. Thus, KO neurons would be expected to exhibit enhanced connectivity at younger ages compared to control cultures. Increased functional connectivity in KO networks, driven by longer dendrites with more synaptic structures likely contributed to the precocious onset of coordinated network bursting behavior observed in MEA experiments. The effects observed on network activity were apparent at the level of individual clones when grouped by *SYNGAP1* genotype. These data further strengthen the conclusion that loss of SynGAP protein drives effects on network activity and these data provide a possible neurobiological mechanism for why individuals with *SYNGAP1* mutations have such a high incidence of early onset pediatric seizures (Vlaskamp et al., 2019).

Data implicating *SYNGAP1* expression on the structural and functional maturation of human neurons is consistent with known functions of this gene discovered from experimentation in mouse neurons (Kilinc et al., 2018). SynGAP protein is highly expressed in rodent neurons and is capable of bidirectional regulation of excitatory synapse strength. Overexpression of SynGAP protein suppresses excitatory synapse transmission by activating AMPA receptor internalization (Rumbaugh et al., 2006). One report indicates that SynGAP isoforms regulate synaptic strength in opposing directions (McMahon et al., 2012). However, genetic ablation of all *Syngap1* splice forms in mice, which removes expression of all protein isoforms, leads to increased excitatory synapse strength and early appearance of synaptic activity in glutamatergic neurons (Clement et al., 2012). These data indicate that the integrated function of all SynGAP proteins in developing mouse neurons is to suppress excitatory synapse function during development. Our findings in human neurons, which also ablated expression of human SynGAP isoforms, support this model of developmental *SYNGAP1* function. Given that we observed early and enhanced excitatory synapse function in KO iNeurons, human *SYNGAP1* also appears to slow the onset of excitatory synapse activity by suppressing excitatory synapse function.

The impact of *SYNGAP1* on human neuron dendritic maturation is also consistent with observations in rodent neurons. The effect of SynGAP protein expression on rodent neuron dendritic development is complex and depends on the type of neuron and brain area studied. *Syngap1* heterozygous KO mice have well documented impairments in dendritic morphogenesis that is linked to alterations in neural circuit assembly and neuronal connectivity. Layer 5 (L5) neurons in the somatosensory cortex of these mutant mice undergo a form of accelerated post-mitotic differentiation, where dendritic extension proceeds at a quicker pace compared to WT mice (Aceti et al., 2015). Interestingly, these neurons also undergo premature spine morphogenesis and early spine pruning. These observations, combined with a desynchronization of L5 cell body and dendritic arbor growth, strongly indicate that SynGAP expression acts in these neurons to suppress a differentiation program that stimulates neuronal maturation. In contrast to these findings, neurons in the upper lamina (Layers 2-4) of the somatosensory cortex of *Syngap1* KO mice show the opposite phenotype. These neurons undergo a form of arrested development where dendritic arbors are shorter compared to similar neurons in WT littermates (Michaelson et al., 2018). Neurons with shorter dendritic arbors also had fewer dendritic spines and these structural alterations impacted connectivity within somatosensory cortex circuits. While our studies in human iNeurons support a role for *SYNGAP1* to suppress dendritic maturation, the specific effect of the gene on structural maturation may also be dependent on the type of human neuron. Two-dimensional neuronal cultures lack the cellular complexity of neural networks found in the intact nervous system. It will be of considerable interest to assess how loss of *SYNGAP1* expression impacts various types of genetically and morphologically distinct neurons formed in three-dimensional human culture systems, such as organoids, and how alterations to dendritic morphogenesis may contribute to impaired neural circuit connectivity and development of network activity.

## Author Contributions

N.L. performed experiments, designed experiments, analyzed data, co-wrote and edited the manuscript. V.A, R. V., M.K., L.B., C.R., A.R., C.H., B.S., and E.W. performed experiments, analyzed data and interpreted data. G.R. conceived project, designed experiments, interpreted data, and co-wrote the manuscript. D.R.P., L.S., T.P.S., C.A.M., and J.L.H. designed and interpreted experiments and edited the manuscript.

## Data and materials availability

hiPSC clones and data supporting the findings of this study are available from the corresponding author upon reasonable request.

